# Sleeve gastrectomy improves metabolic health, cognition, and Alzheimer’s Disease pathology in 3xTG mice

**DOI:** 10.64898/2026.04.21.719988

**Authors:** Reji Babygirija, Julia A. Illiano, Michelle M. Sonsalla, Grace Zhu, Twinkle Mathew, Tristan Molkentin, Meredith Peterson, Carolyn Winder, Fan Xiao, Justin M. Wolter, Dudley W. Lamming, David A. Harris

## Abstract

Obesity and diabetes are well-established risk factors for Alzheimer’s disease (AD), implicating metabolic dysfunction in AD pathogenesis. Sleeve gastrectomy (SG) is among the most effective metabolic interventions available, yet its impact on AD progression remains poorly understood. We hypothesized that SG performed early in life would improve metabolic health, attenuate AD pathology, and preserve cognition in a transgenic AD mouse model. Five-week-old female 3xTg-AD mice were preconditioned on a Western diet (WD) to induce obesity and glucose intolerance, then randomized to SG or sham surgery and maintained on either standard chow or continued WD for 12 months. Metabolic phenotyping, body composition, and cognitive assessments (Novel Object Recognition, Barnes Maze, Fear Conditioning) were performed longitudinally, with histological and molecular analysis of brain tissue at endpoint. Under chow-fed conditions, SG reduced adiposity, improved insulin sensitivity, and decreased cortical Aβ plaque burden, accompanied by attenuated gliosis (GFAP, IBA1) and region-specific, insulin pathway-dependent modulation of autophagy. Under persistent WD, SG improved metabolic health, frailty, spatial cognition, and Aβ pathology, again with corresponding perturbations in autophagy pathway activity. Across all groups, food intake did not differ significantly, indicating that these effects were not secondary to caloric restriction. Collectively, these data suggest that SG engages neuroprotective brain insulin signaling pathways, supporting metabolic surgery as a promising disease-modifying intervention for AD prevention in individuals with obesity.

## Introduction

Alzheimer’s Disease (AD) represents one of the most urgent public health crises of the twenty-first century. As the global population ages, the prevalence of AD is growing at an alarming rate. Approximately 6.9 million Americans aged 65 and older are currently living with the disease, a number projected to nearly double to 13.8 million by 2060 in the absence of any effective preventive or curative treatments (Anon 2024). While recent therapeutic advances mildly impact AD progression, these treatments are associated with significant side effects, high cost, and limited efficacy, highlighting the urgent need for alternative, accessible, and affordable therapeutic strategies for AD.

Metabolic dysfunction is increasingly recognized as a critical contributor to AD pathology. Obesity and diabetes are well-established risk factors for AD, with evidence linking insulin resistance, systemic inflammation, and metabolic dysregulation to neurodegeneration (Khan & Hegde 2020; Pugazhenthi et al. 2017). Beyond cardiovascular complications, accumulating evidence demonstrates that excess adiposity is independently associated with increased risk of dementia and cognitive impairment across the lifespan (Fitzpatrick et al. 2009; Fergenbaum et al. 2009).Targeted dietary interventions including caloric restriction, protein restriction and intermittent fasting have been shown to influence AD progression in mice (Halagappa et al. 2007; Rangan et al. 2022; Babygirija et al. 2024; Babygirija et al. 2026; Babygirija et al. 2025; Kapogiannis et al. 2024). However, the long-term sustainability of such interventions remains a significant challenge as adherence to these dietary regimens are consistently low in humans. Additionally, novel incretin-based therapies targeting the glucagon-like peptide 1 (GLP-1) and gastric inhibitory polypeptide (GIP) receptors have been extensively studied in this context with mixed results in mice (Holscher 2021; Germano et al. 2024; Maskery et al. 2020) and have not been shown to modify disease progression in humans (Cummings et al. 2025; Scheltens et al. 2026).

Among metabolic interventions, bariatric surgery (BS) remains the most effective and durable treatment for substantial, rapid weight loss and resolution of obesity-associated comorbidities (Gloy et al. 2013; Mingrone et al. 2021). Unlike other metabolic interventions mentioned above, bariatric surgery is a one-time intervention with lifelong impact. Sleeve gastrectomy (SG) is now the most performed bariatric procedure worldwide (Gagner et al. 2016). While SG works in part through the induction of incretin hormones to impart metabolic benefits (Chaudhari et al. 2021), incretin signaling is not required to induce weight loss and improvement in diabetes (Wilson-Pérez, Chambers, Ryan, Li, Sandoval, Stoffers, Drucker, Perez-Tilve, et al. 2013), suggesting BS engages a broad array of gut-brain pathways. Furthermore, the benefits of BS extend well beyond metabolic improvements. Neuroimaging studies have demonstrated structural and functional brain changes following BS (Almby et al. 2021; Tuulari et al. 2016; Zhang et al. 2016) and epidemiological data indicate that BS is associated with significant improvements in cognitive function as early as 12 weeks post-operatively, with sustained benefits observed through 24 months (Spitznagel et al. 2013; Gunstad et al. 2011). Remarkably, a recent retrospective cohort study reported that BS was associated with reduced incidence of mild cognitive impairment and AD-related dementias (Chen et al. 2025). Despite these promising findings on cognition and metabolic dysfunction, the impact of BS on AD-specific pathology and the mechanisms driving this neuroprotection remains unknown.

Diet has been identified as one of the key modulators of AD pathology. Long-term consumption of a western diet has been previously shown to exacerbate multiple aspects of AD pathology, including neuroinflammation, β-amyloid pathology, and tau pathology in transgenic mouse models of AD (Sonsalla et al. 2025; Graham et al. 2016; Wiȩckowska-Gacek et al. 2021). Importantly, prior work from our laboratory in C57BL/6J mice demonstrated that a low-protein diet following SG sustained the metabolic benefits of surgery and identified clusters of differentially expressed genes and metabolites associated with the weight-loss phenotype, with effects primarily driven by hepatic tissue (Illiano et al. 2025). However, the downstream effects of SG on AD-related phenotype and cognitive function in the context of diet have not been previously studied.

To address this gap, we used the 3xTg mouse model, which expresses familial human isoforms of APP (APPSwe), Tau (tauP301L), and Presenilin (PS1M146V), and exhibits key features of human AD, including Aβ plaques, neurofibrillary tangles and progressive cognitive deficits (Webster et al. 2014; Oddo et al. 2003). Female 3xTg-AD mice were preconditioned on a western diet (WD) for two months prior to undergoing SG or sham surgery. They were then maintained on one of two lifelong diets: standard chow or WD. This experimental design allowed us to evaluate the effects of SG both under baseline metabolic conditions and in the context of diet-induced metabolic stress.

In the present study, we tested the central hypothesis that SG would attenuate AD progression in 3xTg-AD mice by improving overall metabolic health, reducing neuroinflammation, and decreasing Aβ pathology across dietary contexts. Our results demonstrate that SG reduced adiposity, improved insulin sensitivity, and was associated with reduced cortical amyloid plaque burden and glial activation in chow-fed mice. In mice maintained on a WD, SG had additional benefits including improved spatial learning and memory and reduced plaque pathology. Taken together, these findings provide the first preclinical evidence that SG can slow AD progression in a transgenic AD mouse model, highlighting the potential of metabolic surgery as a disease-modifying intervention for AD and warranting further clinical investigation.

## Results

### SG reduces adiposity and improves insulin sensitivity without altering energy balance in chow-fed 3xTg-AD mice

Five-week-old female 3xTg-AD and Wild-Type (WT) mice were preconditioned on WD (ENVIGO TD.88137) and weight-matched prior to randomization into Sham or SG surgery groups. WD preconditioning was used to induce obesity and glucose intolerance prior to surgery, recapitulating the metabolic comorbidities commonly present in patients seeking bariatric surgery. Following surgery all mice were transitioned back to standard chow diet (TD.180161; 21% protein, 20% fat, 69% carbohydrates by calories) mimicking the post-operative dietary recommendations in clinical practice. Body weight, food intake, and body composition were tracked longitudinally for 12 months post-surgery, with body composition assessed at baseline and endpoint (**Fig.1A).**

**Figure 1:**
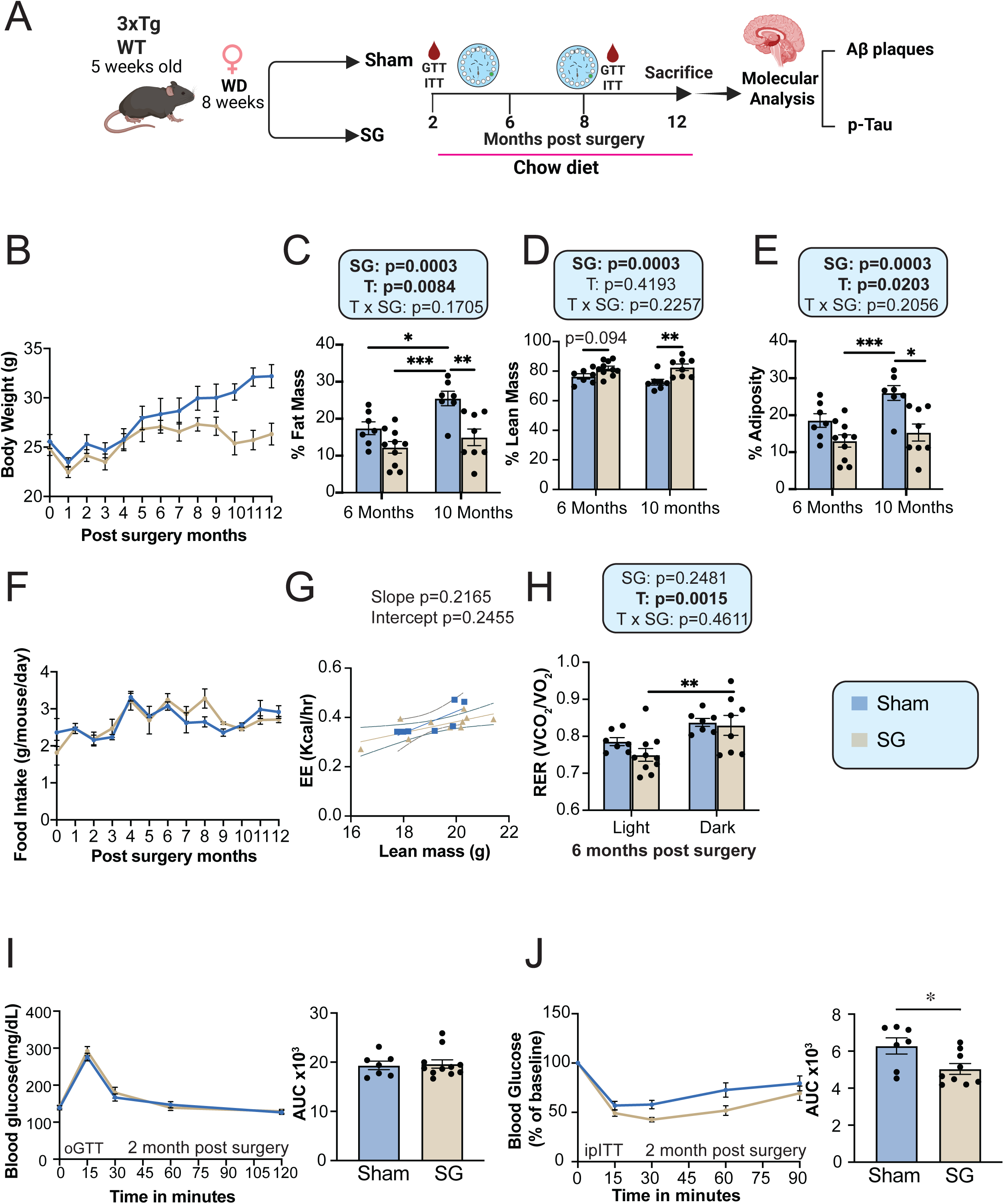
Sleeve gastrectomy reduces adiposity and improves insulin sensitivity independent of food intake in chow-fed 3xTg-AD mice. A) Experimental design of the study using female 3xTg-AD mice. Created with www.biorender.com (B) Bodyweight of female 3xTg-AD mice longitudinally tracked 12 months post-surgery (C) Percent fat mass (D) percent lean mass (E), and percent adiposity assessed by body composition analysis at 6 and 10 months post-surgery. (F) Average daily food intake monitored over 12 months post-surgery. (G) Energy expenditure (EE) during light and dark phases, normalized to body weight at 6 months post-surgery. (H) Respiratory exchange ratio (RER) during light and dark phases at 6 months post-surgery. (I) Oral glucose tolerance test (oGTT) performed at 2 months post-surgery with corresponding area under the curve (AUC). (J) Intraperitoneal insulin tolerance test (ipITT) performed at 2 months post-surgery with corresponding AUC. (B-E). (C-E,G-H) statistics shows the overall effects of surgery (SG), time (T), and their interaction (T×SG) from a two-way ANOVA; with *p<0.05, **p<0.01, ***p<0.001, from Sidak’s multiple comparison test. (I–J) group differences in AUC were assessed by Welch’s t-test; *p<0.05, ***p<0.001. Data represented as mean ± SEM.

As expected, SG 3xTg-AD mice exhibited lower body weight beginning shortly after surgery, which was sustained throughout the 12-month study period (**Fig. 1B**). This was driven primarily by a significant reduction in fat mass at 6 and 10 months post-surgery (**Fig. 1C**), while lean mass was significantly greater in SG mice at both timepoints (**Fig. 1D**). Accordingly, percent adiposity was significantly reduced in SG mice relative to sham controls at both timepoints (**Fig. 1E**). To determine whether reduced food intake accounted for the lower body weight, we examined 12 months of cumulative feeding data. Consistent with our prior work (Harris et al. 2020; Chaudhari et al. 2021; Illiano et al. 2025), caloric intake did not differ between SG and Sham groups (**Fig. 1F**), indicating that reduced caloric intake did not drive the observed differences in body weight and adiposity. WT mice showed analogous reductions in body weight and adiposity without differences in food intake (**Fig. S1A–E**), confirming that the metabolic benefits of SG on body composition are independent of both the AD transgenic background and caloric restriction.

Since SG mice gained less weight than sham mice despite equivalent food intake, we examined energy balance using metabolic chambers. We assessed substrate utilization via the respiratory exchange ratio (RER), calculated as the ratio of CO₂ produced to O₂ consumed. RER approaches 1.0 when carbohydrates are the primary fuel source and 0.7 when lipids predominate. Surprisingly, surgery had no significant effect on energy expenditure (EE) or RER at 6 or 10 months post-surgery (**Fig. 1G–H, S2A-B**). However, a significant effect of time on RER was observed during the dark phase in SG mice at 6 months (**Fig. 1H**), suggesting a transient shift toward greater glucose utilization during the active phase. WT SG mice similarly showed no surgery-driven differences in RER between light and dark phases relative to WT sham controls (**Fig. S1F–G**), indicating this effect is independent of the AD transgenic background. Together, these data demonstrate that reduced adiposity in SG mice is not attributable to differences in total energy expenditure or substrate utilization.

Given that 3xTg-AD mice exhibit progressive impairment in glucose tolerance (Babygirija et al. 2024; Vandal et al. 2015), we examined the impact of SG on glucose handling via an oral glucose tolerance test (oGTT) at two months post-surgery. This revealed no significant differences in glucose excursion or area under the curve (AUC) between SG and Sham groups in either 3xTg or WT mice (**Fig. 1I, S1H**). This finding contrasts with our prior work in non-transgenic C57BL/6J mice, where SG robustly improved glucose tolerance across all dietary conditions (Chaudhari et al. 2021; Harris et al. 2020; Illiano et al. 2025). Nevertheless, an intraperitoneal insulin tolerance test (ITT) at the same timepoint revealed significantly improved insulin sensitivity in 3xTg-SG mice relative to shams, an effect not observed in WT mice (**Fig. 1J, S1I**). These findings suggest that SG improves peripheral insulin sensitivity in 3xTg-AD mice independent of glucose tolerance, likely reflecting compensatory adaptations specific to age and the transgenic background.

Collectively, these data demonstrate that SG reduces adiposity and improves insulin sensitivity in chow-fed female 3xTg-AD mice without altering food intake or energy balance, with the body weight effect mediated primarily through changes in body composition

### SG attenuates AD neuropathology by reducing plaque deposition, tau phosphorylation and gliosis in a brain region-specific manner

Having established that SG improves metabolic health in chow-fed 3xTg-AD mice, we next examined whether these benefits extended to AD neuropathology. We evaluated key pathological hallmarks including Aβ plaque deposition, tau phosphorylation, and gliosis. Brain sections from 15-month-old female 3xTg-AD mice were immunostained for Aβ plaques using 3,3’-diaminobenzidine (DAB) and imaged across multiple magnifications in the cortex and hippocampus (**Fig. 2A**). Quantification revealed a significant reduction in both plaque percent area and plaque number in the cortex of SG mice relative to shams (**Fig. 2B, upper and lower**, respectively). Although there was a trend toward reduced plaque burden observed in the hippocampus (p=0.105), this did not reach statistical significance, indicating that the anti-amyloidogenic effects of SG are cortex-predominant at 15 months We next assessed tau pathology by immunoblotting for phosphorylated tau at threonine 231 (pTau Thr231), a well-established early phosphorylation site in AD-associated tau dysfunction (Alonso et al. 2010) (**Fig. 2C**). Quantification of the pTau/Tau ratio revealed a significant reduction in cortical tau phosphorylation in SG mice relative to shams, with a significant surgery-by-region interaction (p=0.0306), confirming that this effect was cortex-specific with no significant change in hippocampal pTau. The concordance between region-specific plaque and pTau patterns suggests that SG preferentially attenuates early cortical AD pathology in this model.

**Figure 2:**
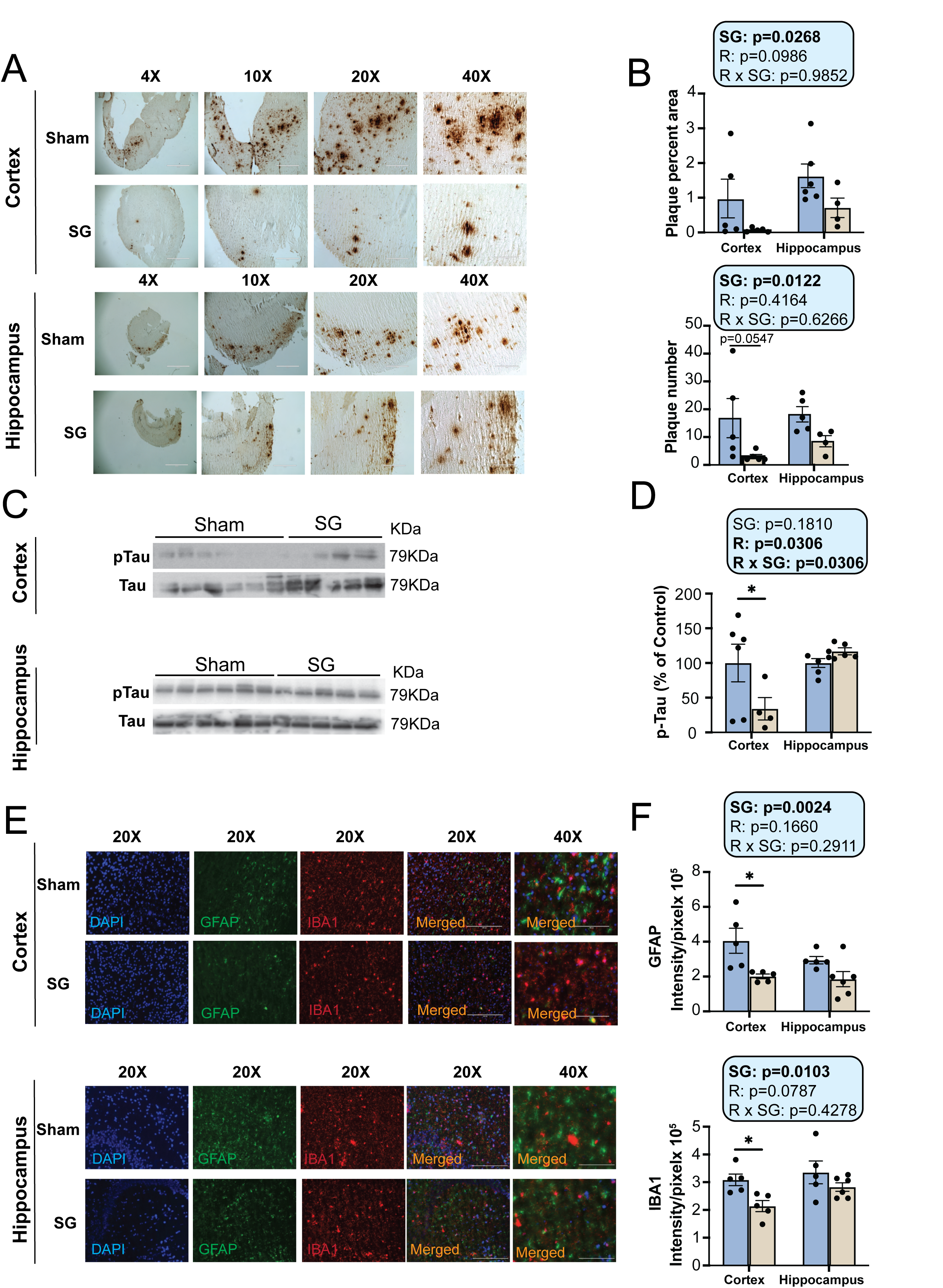
Sleeve gastrectomy reduces cortical amyloid plaque burden, tau phosphorylation, and neuroinflammation in chow-fed 3xTg-AD mice. (A-E) Analysis of AD neuropathology in female 3xTg mice on chow-fed diet for 12 months following SG (A) Representative plaque images of DAB staining with 6E10 antibody in the cortex (top) and hippocampus (bottom) of female 3xTg mice. 4x, 10x, 20x and 40x magnification shown; scale bar in the 4x image is 1000 μM, 10x image is 400μM, 20x is 200 μM and 40x is 100 μM. (B) Quantification of plaque percent area and plaque number in cortex and hippocampus. (C) Representative western blots for phosphorylated tau (pTau Thr231) and total tau in cortex and hippocampus. (D) Quantification of pTau/Tau ratio in cortex and hippocampus. (E) Representative immunofluorescence images of 5 μm paraffin-embedded brain sections co-stained for astrocytes (GFAP) and microglia (IBA1) at 20× magnification. Scale bar = 200 μm. (F) Quantification of GFAP and IBA1 fluorescence intensity in cortex and hippocampus. (B, D, F) statistics shows the overall effects of surgery (SG), brain region (R), and their interaction (R×SG) from a two-way ANOVA; *p<0.05, from Sidak’s multiple comparisons test. An outlier was removed from (D) using ROUT outlier test in Graph Pad prism. Data are represented as mean ± SEM.

Finally, reactive astrogliosis and microglial activation are hallmark features of AD neuroinflammation (Bronzuoli et al. 2019; Adamu et al. 2024; Kiraly et al. 2023). To assess glial responses, brain sections were co-immunostained for glial fibrillary acidic protein (GFAP), a marker of reactive astrocytes, and ionized calcium-binding adaptor molecule 1 (IBA1), a marker of activated microglia (**Fig. 2E**). As expected, given the elevated plaque burden, sham mice exhibited increased GFAP and IBA1 immunoreactivity in both the cortex and hippocampus. In SG mice, both astrocyte activation (GFAP; **Fig. 2F**, upper) and microglial activation (IBA1; **Fig. 2F**, lower) were significantly reduced in the cortex. A trend toward reduced hippocampal glial activation was observed in SG mice (p=0.175) but did not reach statistical significance, consistent with the region-specific pattern of attenuated plaque and tau pathology.

Collectively, these findings demonstrate that SG significantly attenuates multiple hallmarks of AD neuropathology in chow-fed 3xTg-AD mice — including Aβ plaque deposition, tau hyperphosphorylation, and glial activation — with effects predominating in the cortex.

### SG differentially regulates autophagy machinery and insulin signaling in a region-specific manner

Impaired autophagy results in ineffective clearance of plaques and tangles, accelerating disease progression in both patients and animal models (Pickford et al. 2008; Nixon et al. 2005). Given the significant reductions in cortical plaque burden and gliosis observed in SG mice, we investigated whether SG altered autophagy pathway activity in the brain. Cortical and hippocampal lysates from 15-month-old female 3xTg-AD mice were immunoblotted for a panel of autophagy markers: the initiation proteins ATG5, ATG7, and ATG16L1; the autophagosome-associated lipidation marker LC3A/B; and the cargo receptor p62/SQSTM1, whose accumulation inversely reflects autophagic flux. Phosphorylated AKT (pAKT), an upstream inhibitor of autophagy initiation via mechanistic target of rapamycin complex 1 (mTORC1) activation, was also assessed (**Fig. 3A**).

**Figure 3:**
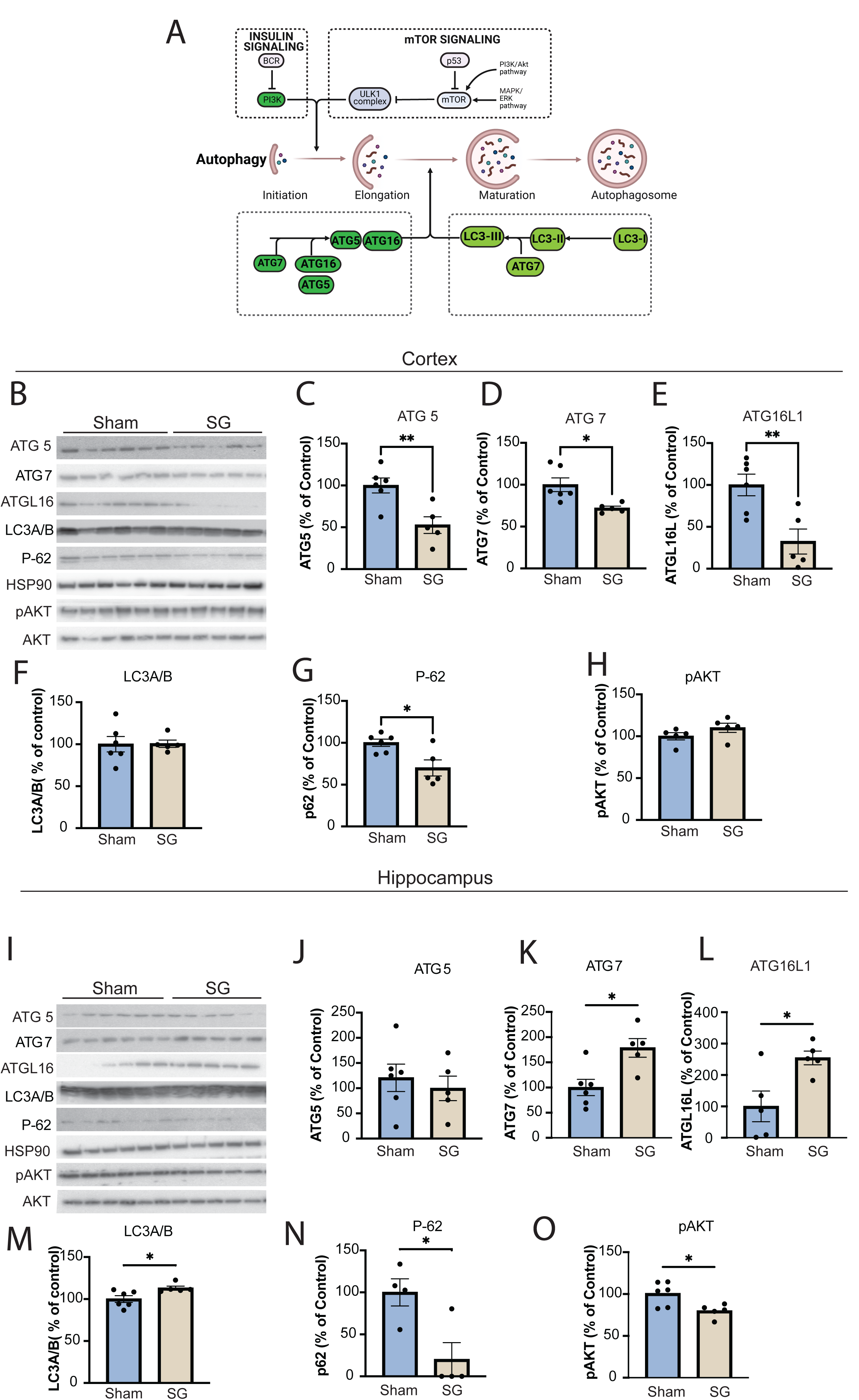
Sleeve gastrectomy differentially regulates autophagy machinery and insulin signaling in a brain-region-specific manner in chow-fed 3xTg-AD mice. (A) Schematic of the autophagy pathway. (B-H) Autophagy marker expression in cortical lysates of 15-month-old female 3xTg-AD mice following SG or sham surgery. (B) Representative immunoblots of the autophagy related proteins in the cortex. Quantification of autophagy proteins (C) ATG5, (D) ATG7, and (E) ATG16L1, as well as autophagosome formation protein (F) light chain 3A/B (LC3A/B), and the autophagy receptor (G) p62 (sequestosome 1, SQSTM1) and (H) pAKT, all normalized to HSP90 expression. (I) Representatives immunoblot of all the autophagy related proteins in Hippocampus. Quantification of ATG5 expression (J), ATG7 (K), ATG16L1 (L), LC3A/B (M), p62 (N) and pAKT (O) relative to expression of HSP90. (C-H, J-O) *p<0.05, **p<0.01, Sham and surgery groups were compared using Welch’s T-test. Data represented as mean ± SEM.

In the cortex, SG mice exhibited significant downregulation of autophagy initiation markers relative to shams (**Fig. 3B**); specifically, ATG5 (**Fig. 3C**), ATG7 (**Fig. 3D**), and ATG16L1 (**Fig. 3E**) were each significantly reduced, indicating suppression of early autophagosome formation. LC3A/B levels were unchanged between groups (**Fig. 3F**), while p62 was significantly reduced (**Fig. 3G**). Rather than reflecting enhanced autophagic flux, the reduction in cortical p62 most likely reflects decreased substrate availability consequent to the significant attenuation of cortical amyloid burden in SG mice. This interpretation is supported by the concurrent downregulation of ATG5, ATG7, and ATG16L1, collectively consistent with a homeostatic reduction in autophagic demand in a region where pathological load has been substantially cleared. A trend toward increased pAKT in SG mice (**Fig. 3H**) further supports suppression of autophagy initiation via mTORC1 activation.

In contrast, the hippocampus displayed a strikingly divergent pattern (**Fig. 3I**). SG mice showed a trend toward increased ATG5 (**Fig. 3J**) with significant upregulation of ATG7 (**Fig. 3K**) and ATG16L1 (**Fig. 3L**), collectively indicating enhanced autophagy initiation. This was accompanied by increased LC3A/B (**Fig. 3M**) and markedly reduced p62 (**Fig. 3N**), indicating augmented autophagosome formation, enhanced flux, and effective cargo clearance. Concordantly, pAKT was significantly reduced in the hippocampus of SG mice (**Fig. 3O**), consistent with relief of mTORC1-mediated autophagy suppression.

Collectively, these data reveal a striking region-specific divergence in autophagy regulation following SG, likely mediated through the AKT/mTORC1 pathway. The trend toward increased cortical pAKT in SG mice is consistent with the improved peripheral insulin sensitivity observed at this timepoint, suggesting that systemic improvements in insulin signaling may extend to the brain and contribute to region-specific modulation of mTORC1-mediated autophagy. The region-specific divergence may also reflect the differential plaque burden across regions. In the cortex, where SG reduced plaque burden, autophagy is correspondingly downregulated, whereas in the hippocampus, where plaque burden remains elevated, autophagy is upregulated, potentially representing a compensatory response to greater pathological load.

### Chow-fed 3xTg AD mice did not exhibit cognitive deficits at 9 or 13 months of age limiting the detection of SG mediated cognition benefits

We next examined whether the neuropathological benefits of SG translated into cognitive improvements by performing a battery of behavioral tests at 5 months (9 months of age) and 8 months (13 months of age) post-surgery. Tests included spatial learning and memory (Barnes Maze; BM), associative fear memory (contextual and cued fear conditioning; FC), and object recognition memory (Novel Object Recognition; NOR). In the BM, mice were trained over four days to locate a hidden escape box using spatial cues, with short-term memory (STM) and long-term memory (LTM) assessed on days 5 and 12, respectively. In FC, mice are placed in a novel chamber and exposed to an auditory tone followed with a 2 second mild foot shock during training. Associative memory was assessed 24 hours later in two phases: contextual fear memory was tested by re-exposing the mice to original training chamber in the absence of tone, and cued memory test was assessed in a novel context with tone presentation. Freezing behavior was recorded as the measure of fear memory in both these phases. In the NOR test, which exploits a mouse’s natural preference for novelty, exploration of novel and familiar objects was quantified via the discrimination index (DI) — calculated as the difference in time exploring the novel versus familiar object divided by total exploration time — at both STM and LTM recall. A positive DI indicates preferential exploration of the novel object and is interpreted as consolidated memory.

At 5 months post-surgery (9 months of age), SG and sham mice showed comparable acquisition of the escape hole location across the four-day BM training phase, with no significant differences in latency to goal or error hole visits at either STM or LTM (**Fig. 4A–B**). Importantly the sham 3xTg mice performed comparably to WT controls across all behavioral parameters at this time point **(Figs. S3A-S3E)**. In the FC paradigm, no significant surgery-by-group interactions were detected during cued (**Fig. 4C**) or contextual acclimation phases (**Fig. 4D**). However, there was a significant effect of time across trial phases reflecting normal learning in both groups (**Fig. 4C–D**). In NOR testing, neither group showed a significant preference for the novel object at STM or LTM, and DI and absolute DI values did not differ between groups (**Fig. 4E–F**).

**Figure 4:**
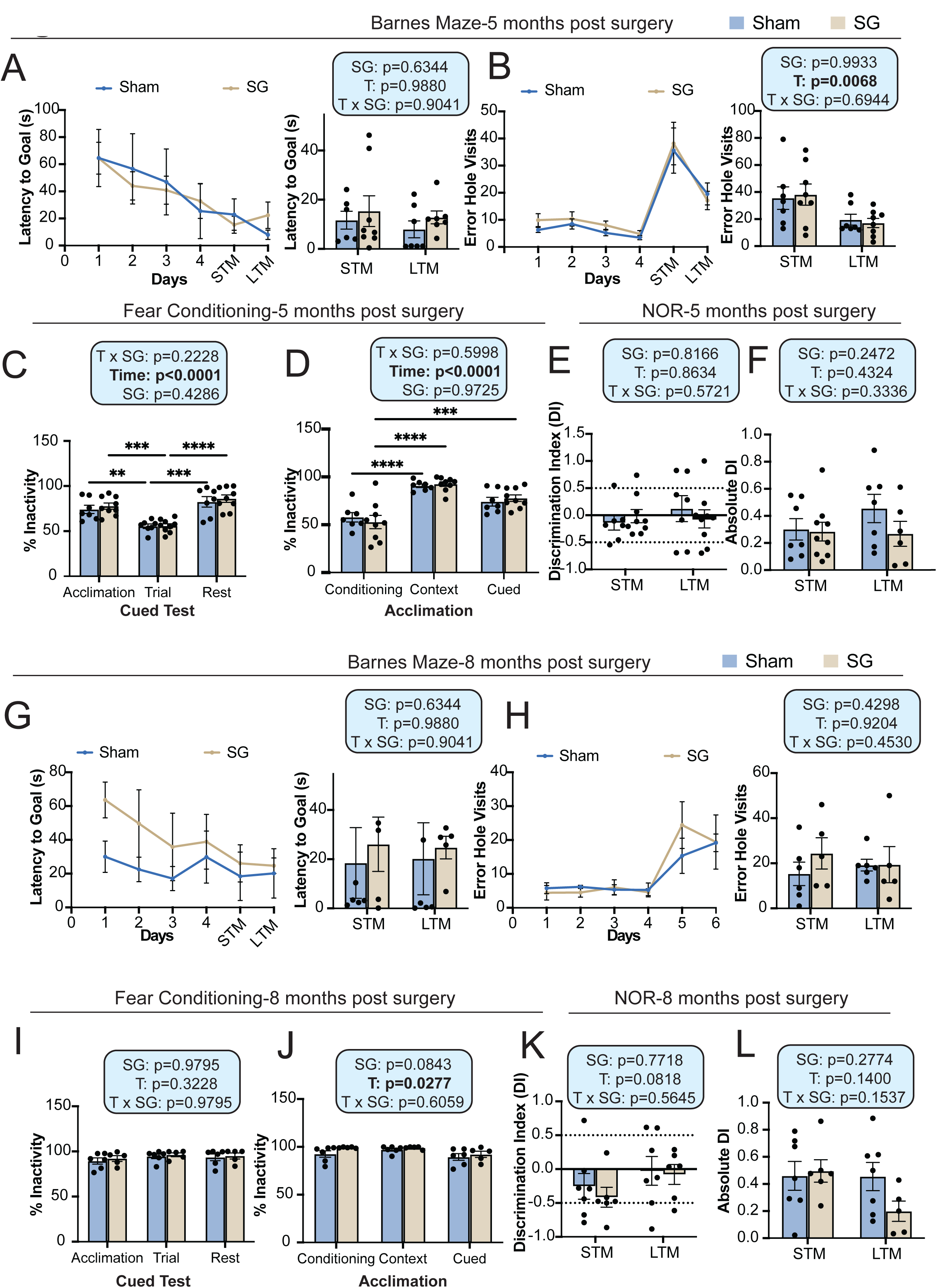
Chow-fed 3xTg-AD mice do not exhibit cognitive deficits up to 8 months post-surgery. (A, G) Latency of target in Barnes Maze acquisition period over the five days of training and in short term memory (STM) and long-term memory (LTM) tests at 5 months (A) and 8 months (G) post-surgery. (B, H) The number of error hole visits during Barnes maze training phase in STM and LTM tests at 5 months (B) and 8 months (H) post-surgery. (C, I) Percent inactivity recorded during the cued test phase of the fear conditioning paradigm at 5 months (C) and 8 months (I) post-surgery. (D, J) Percent inactivity recorded during the acclimation, conditioning, context, and cued phases of the fear conditioning paradigm at 5 months (D) and 8 months (J) post-surgery (E, K) The preference for a novel object over a familiar object was assayed at 5 months (E) and 8 months post-surgery (K) via STM and LTM tests. The dashed lines at +0.2 and -0.2 indicates the threshold for discrimination index (DI) values showing the preference for novel or familiar objects. (F, L) Absolute DI was plotted to show the magnitude of discrimination regardless of the direction of preference at 5 months (F) and 8 months (L) post-surgery in 3xTg-AD mice. (A-L) Statistics for the overall effects of surgery, time and the interaction represent the p value from a 2-way ANOVA. **p<0.01, ***p<0.001 from Sidak’s multiple comparison test.

Behavioral testing repeated at 8 months post-surgery (13 months of age) again revealed no significant group differences across any cognitive domain. BM performance at STM and LTM did not differ between SG and sham mice in latency to goal or error hole visits (**Fig. 4G–H**), FC was unaffected by surgery during cued (**Fig. 4I**) and contextual acclimation phases (**Fig. 4J**), and NOR discrimination indices and absolute DI values at STM and LTM did not differ between groups (**Fig. 4K–L**) The absence of detectable cognitive differences at both timepoints most likely reflects insufficient cognitive deficit in chow-fed 3xTg-AD mice at these ages under the tested conditions, rather than a true absence of SG effect. To address this, we examined the impact of SG on metabolism, cognition, and AD-specific pathology under lifelong WD exposure. Long-term WD consumption exacerbates multiple hallmarks of AD pathology — including neuroinflammation, Aβ deposition, and tau pathology — in transgenic mouse models (Wiȩckowska-Gacek et al. 2021; Graham et al. 2016; Sonsalla et al. 2025), providing a disease context in which the compound effects of genetic and dietary risk factors interact to exacerbate AD outcomes. WD represents a clinically relevant metabolic stressor which when combined with AD-prone genetics can recapitulate the metabolic burden present in the human disease spectrum. We hypothesized that the combined metabolic and neuropathological burden of persistent WD feeding would unmask pro-cognitive benefits of SG not detectable in the chow-fed cohort.

### SG reduces adiposity, shifts substrate utilization, and transiently restores glucose homeostasis in 3xTg-AD mice under persistent western diet conditions

A separate cohort of female 3xTg-AD mice was preconditioned on WD for 8 weeks prior to surgery and maintained on WD throughout the 12-month post-surgical period (**Fig. 5A**). Body weight, food intake, body composition, and metabolic parameters were monitored longitudinally, with glucose homeostasis assessed at 1 and 9 months post-surgery. Consistent with the chow-fed cohort, SG mice showed attenuated weight gain relative to shams throughout the study period (**Fig. 5B**), driven by a significant reduction in percent fat mass at 5 and 8 months post-surgery (**Fig. 5C**) and a trend toward greater lean mass (**Fig. 5D**). Food intake did not differ significantly between groups (**Fig. 5E**), confirming that reduced caloric intake did not account for the lower adiposity observed in SG mice, regardless of dietary context.

**Figure 5:**
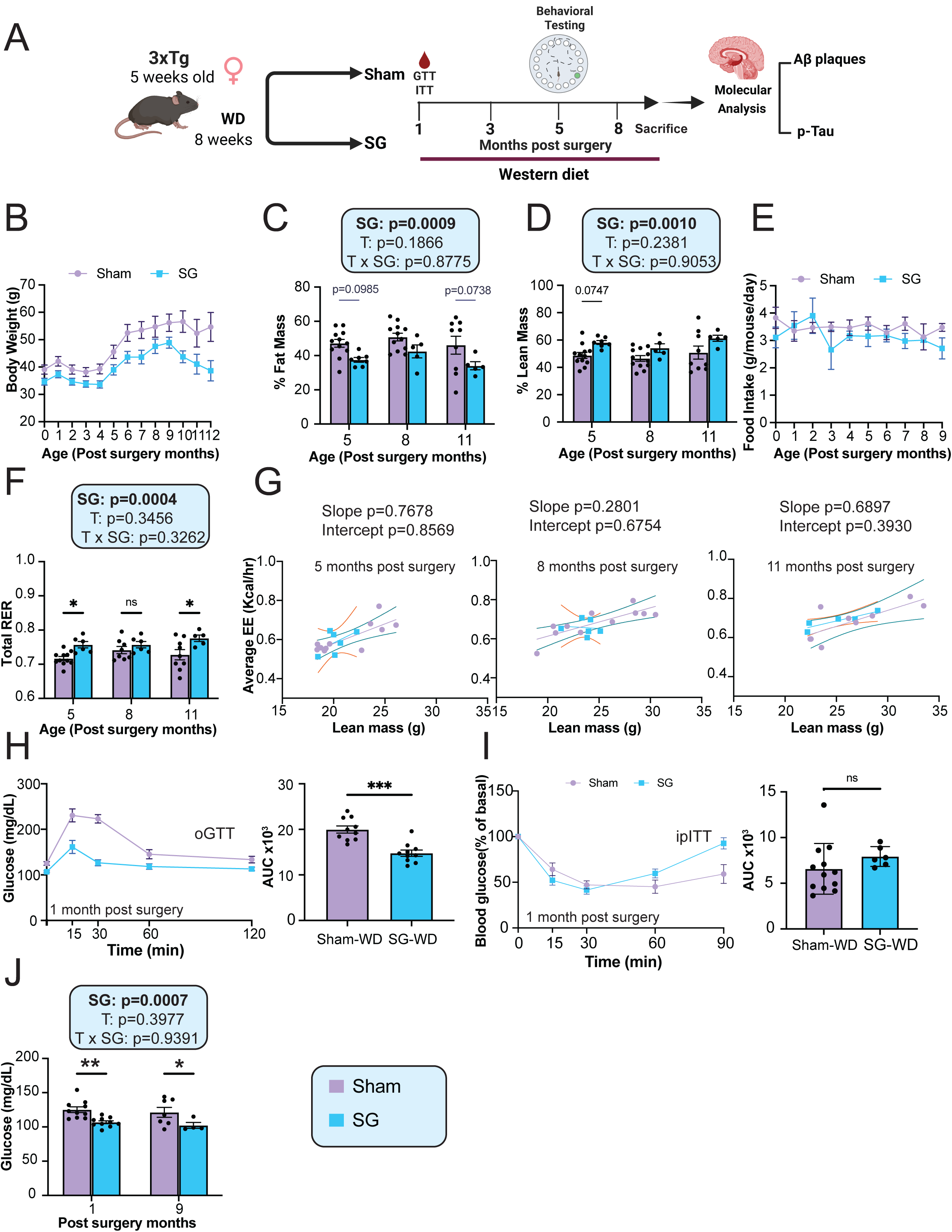
Sleeve gastrectomy reduces adiposity and improves glucose tolerance in WD-fed 3xTg-AD mice. A) Experimental design of the study using female 3xTg-AD mice (B) Bodyweight of female 3xTg-AD mice longitudinally tracked 12 months post-surgery (C) Percent fat mass, (D) Percent lean mass assessed by body composition analysis at 5, 8 and 11 months post-surgery. (E) Average daily food intake monitored over 12 months post-surgery. (F) Respiratory exchange ratio (RER) during light and dark phases at 5,8 and 11 months post-surgery. (G) Energy expenditure (EE) during light and dark phases, normalized to body weight at 5, 8 and 11 months post-surgery. (H) Oral glucose tolerance test (oGTT) performed at 1-month post-surgery with corresponding area under the curve (AUC). (I) Intraperitoneal insulin tolerance test (ipITT) performed at 1-month post-surgery with corresponding AUC. (B-E). (J) Fasting blood glucose levels at 1 and 9 months post-surgery in female 3xTg AD mice. (C-D,F-G,J) statistics shows the overall effects of surgery (SG), time (T), and their interaction (T×SG) from a two-way ANOVA; with *p<0.05, **p<0.01, ***p<0.001, from Sidak’s multiple comparison test. (H, I) group differences in AUC were assessed by Welch’s t-test; *p<0.05, **p<0.01, ***p<0.001. Data represented as mean ± SEM. (A) Created with www.Biorender.com

To assess substrate utilization, mice were placed in metabolic chambers. SG mice exhibited a significantly higher RER than shams across the study period (**Fig. 5F**), with values consistently above 0.7, reflecting a shift toward mixed fuel substrate utilization, which was most pronounced at 5 and 11 months post-surgery. ANCOVA with lean mass as a covariate revealed no significant differences in slopes or elevations between groups at 5, 8, or 11 months post-surgery, indicating that energy expenditure was comparable between groups after accounting for lean mass (**Fig. 5G**)

An oGTT at 1-month post-surgery demonstrated that WD-fed sham mice were glucose intolerant, while SG mice showed fully restored glucose tolerance (**Fig. 5H**), indicating that SG rapidly reversed diet-induced glucose intolerance. An ipITT at the same timepoint revealed no significant difference in insulin sensitivity between groups (**Fig. 5I**), suggesting that the improvement in glucose tolerance occurred independently of peripheral insulin action. Notably, fasting blood glucose was significantly reduced in SG mice at both 1 and 9 months post-surgery under persistent WD conditions (**Fig. 5J**). By 9 months, however, glucose tolerance and insulin sensitivity no longer differed between groups (**Fig. S4A–B**), driven largely by progressive normalization of glucose tolerance in the sham cohort over time.

Collectively, these data demonstrate that SG confers meaningful metabolic protection in female 3xTg-AD mice under persistent dietary stress, reducing adiposity, shifting substrate utilization toward carbohydrate oxidation, and lowering fasting glycemia — broadly consistent with the metabolic benefits observed in the chow-fed cohort, though with a distinct effect on substrate utilization not seen under standard dietary conditions.

### SG attenuates AD neuropathology, rescues cognitive deficits, and reduces frailty progression under persistent western diet conditions

We assessed spatial learning and memory using the Barnes Maze at 9 months post-surgery and elected not to perform NOR or FC testing to reduce stress on the remaining animals. In contrast to the chow-fed cohort where no group differences were detected (**Fig. 4**), WD-fed Sham mice exhibited LTM tests compared to SG mice (**Fig. 6A, left).** Consistent with this, SG mice demonstrated a significantly higher success rate in locating the escape box at both STM and LTM compared to sham-WD controls (**Fig. 6A, right)**.

**Figure 6:**
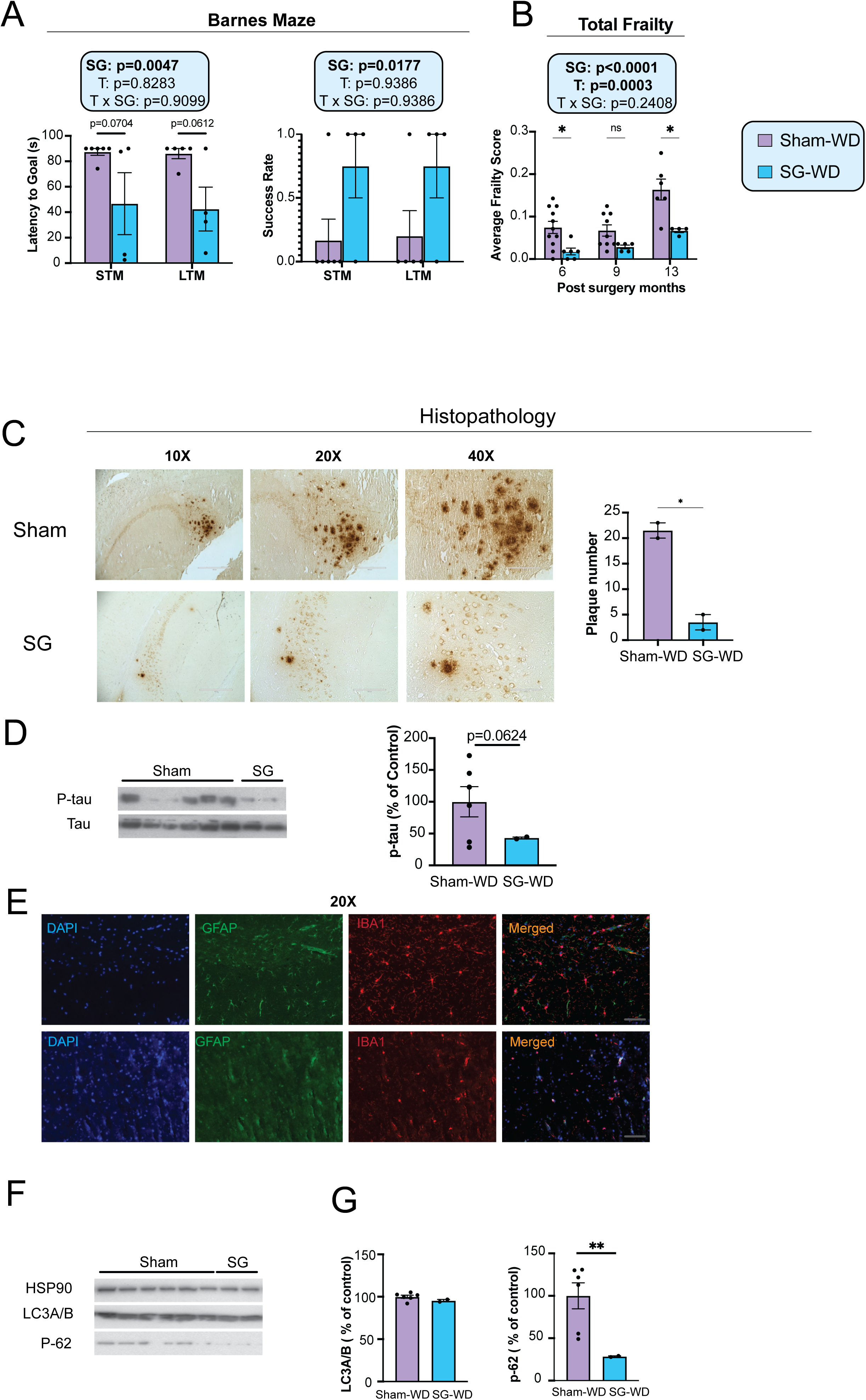
Sleeve gastrectomy rescues spatial memory deficits, reduces frailty, and improves AD-specific pathology in Western diet-fed 3xTg-AD mice. (A) Latency to the target hole and success rate during STM and LTM probe trials of the Barnes Maze test at 9 months post-surgery. (B) Average frailty index scores assessed longitudinally at 6, 9, and 13 months post-surgery. (C) Representative plaque images of DAB staining with 6E10 antibody in the brains of female 3xTg-AD mice. 4x, 10x, 20x and 40x magnification shown; scale bar in the 4x image is 1000 μM, 10x image is 400μM, 20x is 200 μM and 40x is 100 μM. Quantification of plaque number. (D) Representative western blots for phosphorylated tau (pTau Thr231) and total tau and the quantification of pTau/Tau ratio. (E) Representative immunofluorescence images of 5 μm paraffin-embedded brain sections co-stained for astrocytes (GFAP) and microglia (IBA1) at 20× magnification. Scale bar = 200 μm. (F) Representative immunoblots and quantification of LC3A/B and p62/SQSTM1 in brain lysates, normalized to HSP90. (G) Quantification of LC3A/B and p62 relative to expression of HSP90. For panels C, D and G, groups were compared using Welch’s t-test; *p<0.05. For panels A and B, the overall effects of surgery (SG), time (T), and their interaction (T×SG) were assessed by two-way ANOVA; *p<0.05. Data are represented as mean ± SEM.

Beyond cognitive outcomes, we assessed overall healthspan using a validated mouse frailty index based on the accumulation of age-associated deficits (Whitehead et al. 2014). As expected, frailty scores increased with age in both groups (**Fig. 6B).** Most importantly, SG mice exhibited significantly lower frailty scores than sham-WD controls across the duration of the study (**Fig. 6B).**

The health burden of lifelong WD on this mouse cohort was reflected in the unexpectedly high mortality across both sham and SG groups through the study duration. This unfortunately substantially reduced our final group sizes. Despite this limitation, neuropathological assessment of brain tissue from all surviving mice revealed compelling differences between groups. DAB immunostaining of brain sections from 15-month-old, WD fed 3xTg-AD mice showed a striking and significant reduction in plaque number in SG mice compared to sham-WD controls (**Fig. 6C**) consistent with the anti-amyloidogenic effects observed in the chow-fed cohort (**Fig. 2).** This was accompanied by a reduction in p-Tau levels in SG mice, though given the reduced sample size this did not reach statistical significance (**Fig. 6D**). Immunofluorescence co-staining for GFAP and IBA1 demonstrated visually reduced astrocytic and microglial activation in SG mice relative to Sham-WD controls (**Fig. 6E),** indicating attenuation of neuroinflammation consistent with the reduced plaque burden. Formal quantification was limited by available sample numbers. We also examined autophagy pathway markers in WD-fed mice. Immunoblotting for LC3A/B and p62 revealed that LC3A/B levels did not differ significantly between SG and sham-WD groups (**Fig. 6F-G, left)**, while p62 was significantly reduced in SG mice (**Fig. 6G, right).**

Taken together, we demonstrate that SG significantly attenuates AD neuropathology, rescues WD-accelerated spatial memory deficits, and reduces frailty progression in 3xTg-AD mice maintained on WD.

## Discussion

Type 2 diabetes and obesity are well-established risk factors for AD (Silva et al. 2019). Although the precise mechanistic relationship remains incompletely understood, glucose metabolism is impaired in AD, with documented defects in glucose uptake (Duran-Aniotz & Hetz 2016) and mitochondrial function (Baek et al. 2017; Newington et al. 2013; Krako et al. 2013). Obesity confers a nearly 2-fold increase in AD risk relative to normal-weight individuals (Whitmer et al. 2007), with obesity-driven metabolic dysfunction promoting neuroinflammation and neuronal damage through mechanisms that accelerate disease progression (Valcarcel-Ares et al. 2019; Tarantini et al. 2018; Tucsek et al. 2014). Diet further modulates AD risk, as saturated fats and refined carbohydrates, the principal components of a Western diet, promote excess weight gain and are associated with increased incidence of AD and cognitive dysfunction (Clemente-Suárez et al. 2023; Hu et al. 2001; Valcarcel-Ares et al. 2019).

Considerable effort has been directed toward understanding how targeted metabolic interventions influence AD progression. Dietary and behavioral strategies — including caloric restriction (Halagappa et al. 2007; Rangan et al. 2022), protein restriction (Babygirija et al. 2024), branched-chain amino acid restriction (Babygirija et al. 2026), and intermittent fasting (Babygirija et al. 2025; Kapogiannis et al. 2024) — have each demonstrated impact on AD pathogenesis in transgenic mouse models. However, these approaches are poorly sustained in humans, where attrition rates in dietary and behavioral programs are exceedingly (Fildes et al. 2015; Volkmar et al. 1981; Ponzo et al. 2021). Pharmacological approaches targeting the incretin axis have similarly shown mixed efficacy in preclinical AD models (Holscher 2021; Germano et al. 2024; Maskery et al. 2020) and failed to demonstrate cognitive benefit in clinical trials (Cummings et al. 2026). Furthermore, despite strong efficacy in controlled trial settings, real-world use of GLP-1 and GIP agonists is associated with substantially reduced effectiveness (Powell et al. 2023; Kelly et al. 2024), high therapy attrition (Miller & Brennan 2015), and weight recurrence following cessation (Horn et al. 2026; Wilding et al. 2022).

Multiple trials have established bariatric surgery as the most effective intervention for obesity and obesity-associated metabolic disease, with SG achieving rapid, durable weight loss and meaningful improvement in glycemic control and overall metabolic health (Surgery 2021; Schauer et al. 2017; Heshmati et al. 2019; Mingrone et al. 2015). Critically, as a single intervention, bariatric surgery confers long-term benefits across multiple organ systems (Maciejewski et al. 2016; Syn et al. 2021; Shikora et al. 2022). Clinical evidence regarding the cognitive effects of bariatric surgery, however, remains mixed. Most studies report meaningful cognitive improvements within weeks of surgery and sustained benefits through 6 months post-SG (Tucker et al. 2020), with one recent retrospective cohort study reporting a reduced incidence of mild cognitive impairment and AD-related dementias following surgery (Chen et al. 2025). In contrast, a large cohort study of over 51,000 subjects reported increased dementia incidence in the surgical group over a 10.5-year follow-up period (Kim et al. 2023).

The preclinical study of bariatric surgery on AD has been limited. In a study of Roux-en-Y gastric bypass (alternative form of bariatric surgery), male APP/PS1/TAU mice had improved cognition, downregulated Tau phosphorylation, and reduced Aβ deposition. This was associated with upregulated hippocampal GLP-1, which is associated with improvements in AD (Mei et al. 2023). Notably, however, those animals were not obese, which limits the translational relevance to the clinical population most likely to undergo bariatric surgery. To our knowledge, no prior study has examined the effects of SG specifically on AD pathology progression, which is a critical gap given the well-established links between metabolic dysfunction and neurodegeneration. The present study addresses this gap by characterizing the effects of SG on AD-related phenotype in a well-validated transgenic mouse model The metabolic benefits of SG observed here are consistent with our prior work (Illiano et al. 2025). Under both chow and WD conditions, SG mice maintained lower body weight driven primarily by reduced fat mass accumulation, with no corresponding differences in food intake. Thus, indicating that metabolic improvements were mediated through changes in body composition rather than caloric restriction. These findings extend our previous observations of robust SG-driven metabolic improvements in C57BL/6J mice across multiple dietary conditions (Chaudhari et al. 2021; Harris et al. 2020; Illiano et al. 2025) to a transgenic AD background, confirming that the metabolic efficacy of SG is preserved in the context of AD-related phenotype.

Most importantly, these metabolic improvements translated to meaningful attenuation of AD neuropathology, most strikingly evidenced by a significant reduction in plaque burden selectively in the cortex. Concordantly, cortical pTau was significantly reduced while hippocampal pTau remained unchanged. This region-specific pattern is consistent with the well-described spatiotemporal progression of AD, in which plaque deposition first emerges in neocortical regions before spreading to the hippocampus (Selkoe 1991; Hyman et al. 1984). This preferential cortical attenuation suggests that SG may intervene at an early stage of pathological propagation, prior to hippocampal involvement or alternatively induce cortical-plaque removal. Collectively, the reduction in cortical amyloid burden, tau phosphorylation, and neuroinflammation provides compelling preclinical evidence that SG is neuroprotective, and further investigation into the precise mechanisms driving these effects is warranted.

This is consistent with a region-specific change in the insulin–autophagy axis and active autophagic clearance in SG mice and suggests that restoration of insulin sensitivity may be a key upstream driver of the neuroprotective effects observed in the 3xTg-AD model.

Under chow-fed AD conditions, SG produced a significant improvement in peripheral insulin sensitivity, which is well-documented following SG in both humans and mice (Chaudhari et al. 2021; CĂTOI et al. 2016; Soares et al. 2023; Abu-Gazala et al. 2018). This is particularly relevant given the well-established role of brain insulin signaling in AD pathogenesis (Pliszka & Szablewski 2026; Liu et al. 2011). Cerebral insulin resistance impairs the PI3K/AKT/mTOR signaling axis, leading to dysregulation of autophagy and impaired clearance of Aβ and hyperphosphorylated tau (Razani et al. 2021; Ebrahim et al. 2024). Restored peripheral insulin sensitivity following SG may therefore improve central insulin signaling, promoting region-specific modulation of autophagic flux and plaque clearance. Consistent with this interpretation, pAKT levels were inversely associated with autophagy pathway activation across brain regions. In the cortex, SG mice exhibited elevated pAKT alongside downregulation of ATG5, ATG7, and ATG16L1 indicative of inactivation of the autophagy pathway in a region where plaque burden is already reduced. In contrast, in the hippocampus SG mice showed reduced pAKT, upregulation of the same initiation markers, and decreased p62, consistent with autophagy activation, enhanced autophagic flux and active cargo clearance in a region which has greater residual pathological burden. Together, these findings are consistent with a region-specific shift in the insulin–autophagy axis and suggest that restoration of peripheral insulin sensitivity may be a key upstream driver of the neuroprotective effects observed in this model A notable finding was the dissociation between neuropathological improvement and cognitive outcomes in chow-fed 3xTg-AD mice. Despite significant reductions in plaque burden, tau phosphorylation, and neuroinflammation, there were no detectable improvements in memory or cognitive performance across our testing battery at 5 or 8 months post-surgery. However, this likely reflects insufficient baseline cognitive deficit in the sham-operated chow-fed cohort at these time points rather than a true absence of pro-cognitive effect. In essence, the sham mice were uncharacteristically smart thus precluding detection of SG-mediated cognitive benefits. This interpretation is consistent with the well-documented phenotypic variability of the 3xTg-AD model, in which the onset and severity of cognitive deficits and pathological progression vary considerably across studies and institutions (Javonillo et al. 2022; Oddo et al. 2003; Sterniczuk et al. 2010; Babygirija et al. 2026; Babygirija et al. 2025).

Under persistent dietary stress, sham animals exhibited notably worse metabolic health and accelerated cognitive decline, providing sufficient deficit range to unmask SG-mediated cognitive benefits. The absence of cognitive improvement in the chow-fed cohort therefore reflects a limitation of the experimental model under standard dietary conditions rather than a true absence of SG effect on brain function.

A limitation of the WD arm was reduced cohort size due to age-associated mortality across multiple experimental groups, though final numbers remained broadly consistent with those reported in other long-term 3xTg-AD studies (Babygirija et al. 2024; Babygirija et al. 2025; Julien et al. 2010). Nevertheless, surviving SG mice demonstrated robust and consistent benefits across metabolic, neuropathological, cognitive, and physical health domains including improved glucose homeostasis, reduced amyloid and tau pathology, attenuated neuroinflammation, rescued spatial memory, and reduced frailty. This collectively supports the conclusion that SG attenuates AD progression even under sustained metabolic stress. While replication in larger cohorts is necessary, these findings provide strong preclinical proof-of-concept for SG as a disease-modifying intervention in obesity-associated AD.

Several limitations of the present study warrant consideration. First, we exclusively used female 3xTg-AD mice. Inclusion of male mice and additional AD mouse models would provide broader insight into how SG-mediated benefits generalize across sexes and genetic backgrounds. Male 3xTg-AD mice were not included as they do not reliably develop AD-relevant phenotype in our hands and suffer premature mortality. Second, tau phosphorylation was assessed at a single site, Thr231, a well-established early and clinically relevant phosphorylation event in AD (Ashton et al. 2021). However, multiple sites contribute to neurofibrillary tangle formation and disease progression. The effect of SG on additional phosphorylation sites, including Ser202, Thr205, and Ser396, remains to be examined. Third, while our autophagy analyses revealed region-specific differences in ATG proteins, LC3A/B, and p62, we did not directly measure autophagic flux, which would be required to definitively distinguish changes in autophagy initiation from impaired downstream degradation. Future studies employing flux assays or lysosomal inhibitors would clarify the functional significance of the observed protein-level changes. Fourth, we did not directly measure brain insulin sensitivity, and we only infer these findings from peripheral assays. Fifth, while our IHC and molecular analysis distinguished cortical and hippocampal regions broadly, we did not perform sub-region-specific analysis within these regions. Future studies examining discrete sub regions-such as CA1, CA3 and the dentate gyrus (DG) of hippocampus, or prefrontal, entorhinal cortices would provide greater spatial resolution along with region specific heterogeneous responses to SG that are hidden in bulk tissue analysis. Finally, molecular analyses were conducted on bulk tissue lysates from discrete brain regions. A more spatially resolved approach — single-cell or region-specific transcriptomics and proteomics — will be necessary to fully delineate the mechanistic pathways through which SG exerts neuroprotective effects across distinct cell types and neural circuits.

In summary, this study provides the first preclinical evidence that SG attenuates AD progression in a transgenic mouse model, with benefits spanning metabolic, neuropathological, and cognitive domains. SG reduced adiposity and improved insulin sensitivity across both dietary conditions, and these systemic improvements were accompanied by region-specific reductions in cortical amyloid plaque burden, tau phosphorylation, and glial activation. Cognitive benefits were observed specifically in the WD cohort, where diet-exacerbated pathology generated sufficient behavioral impairment to unmask the pro-cognitive potential of SG. Region-specific modulation of autophagy machinery and insulin signaling further suggests that SG engages distinct neuroprotective mechanisms across cortical and hippocampal circuits Critically, these neuroprotective effects were achieved through a single surgical intervention rather than sustained dietary compliance, distinguishing SG from caloric restriction and other fasting-based approaches that are undermined by poor long-term adherence and weight recurrence (Fildes et al. 2015; Horn et al. 2026; Goudswaard et al. 2023). Moreover, the observed benefits are unlikely to be mediated solely through the GLP-1 axis, given that the metabolic effects of SG are preserved in GLP-1 receptor knockout mice (Wilson-Pérez, Chambers, Ryan, Li, Sandoval, Stoffers, Drucker, Pérez-Tilve, et al. 2013), indicating that SG engages broader and more durable metabolic reprogramming than GLP-1 receptor agonism alone. This mechanistic breadth may underlie the efficacy of SG in a context where pharmacological GLP-1-based therapies have failed to demonstrate cognitive benefit in trial (Cummings et al. 2026). Collectively, these findings support SG as a candidate disease-modifying intervention for patients with obesity and diet-induced metabolic dysfunction prone to AD. Determining the mechanisms through which SG modulates AD risk may inform the development of novel therapeutic strategies for AD prevention.

## Materials and Methods

### Animals

Male and female homozygous 3xTg-AD mice were obtained from The Jackson Laboratory (Bar Harbor, ME, USA) and were bred and maintained at the vivarium with food and water available ad libitum. At five weeks of age, they were preconditioned on a high-fat WD (Envigo, TD.88137; 42% calories from fat; 15% calories from protein; 43% calories from carbohydrates) for 8 weeks to induce obesity and glucose intolerance. Male mice were not used during this study and thus the impact of sex was not studied. Mice were housed in groups of two and had ad libitum access to food and water in a climate-controlled environment with a 12-h light/dark cycle. All procedures were approved by the Institutional Animal Care and Use Committee (UW-Madison and William S. Middleton Memorial Veteran’s Hospital), and animals were cared for according to guidelines set forth by the American Association for Laboratory Animal Science in an AALAC-accredited facility.

### Sleeve gastrectomy and sham procedures

At 12 weeks of age, mice were weight matched and randomized into either a Sham or SG surgical group. In brief, mice were placed on a recovery gel diet from 48 h prior to surgery to 6 days post-surgery before going on their respective diets (Clear H20, Westbrook, ME). Mice were anesthetized with isoflurane under sterile conditions. At the time of surgery, mice received weight-based saline, norocillin, and buprenorphine (Ethiqua XR; Fidelis Animal Health, North Brunswick, NJ). SG consisted of a midline laparotomy, short vessel ligation, and removal of roughly 75% of the stomach and the entirety non-glandular stomach using a linear cutting stapler (Medtronic, Minneapolis, MN). Sham surgery similarly consisted of a laparotomy, short vessel ligation, and manipulation of the stomach along the would-be staple line.

### Glucose and Insulin tolerance tests

OGTT and ITT were performed after a four-hour fast (7 a.m.–11 a.m.). Mice received an oral bolus of 30% glucose (1g/kg) and blood glucose was measured at time 0, 15, 30, 60, and 120 min. For ITT mice received an intraperitoneal bolus of regular insulin (0.5 u/kg; Humulin R: Eli Lilly USA). Blood glucose was measured 0, 15, 30, 60, and 90 min. The blood glucose measurements were collected using a Bayer Contour glucose meter and test strips.

### Body composition and indirect calorimetry analysis

Body composition was assessed using the EchoMRI Body Composition Analyzer. Indirect calorimetry was performed using Oxymax/CLAMs metabolic chamber system. Mice were individually housed in CLAMs cages at 22°C and measurements for CO2 production, O2 consumption, heat production, RER and food intake were recorded for each mouse. Mice were placed in CLAMs cages for ∼48 h with a 12-h light and dark cycle. The first 24 h were considered the acclimation period and were discarded from the analysis. Mice had access to food and water ad libitum.

### Behavioral assays

All mice underwent behavioral phenotyping at 12 months of age. The Novel object recognition test (NOR) was performed in an open field where the movements of the mouse were recorded via a camera that is mounted above the field. Before each test mice were acclimatized in the behavioral room for 30 minutes and were given a 5-minute habituation trial with no objects on the field. This was followed by test phases that consisted of two trials that were 24 hours apart: Short term memory test (STM and Long-term memory test (LTM). In the first trial, the mice were allowed to explore two identical objects placed diagonally on opposite corners of the field for 5 minutes. Following an hour after the acquisition phase, STM was performed and 24 hours later, LTM was done by replacing one of the identical objects with a novel object. The results were quantified using a discrimination index (DI), representing the duration of exploration for the novel object compared to the old object.

For Barnes maze, the test involves 3 phases: habituation, acquisition training and the memory test. During habituation, mice were placed in the arena and allowed to freely explore the escape hole, escape box, and the adjacent area for 2 minutes. Following that during acquisition training the mice were given 180 seconds to find the escape hole, and if they failed to enter the escape box within that time, they were led to the escape hole. After 4 days of training, on the 5th day (STM) and 12th day (LTM) the mice were given 90 second memory probe trials. The latency to enter the escape hole, distance traveled, and average speed were analyzed using Ethovision XT (Noldus).

Associative fear memory was assessed using a standard cued and contextual fear conditioning paradigm. On the conditioning day, mice were placed in the fear-conditioning chamber (26 × 26 × 24 cm) and allowed an 8-minute habituation period, during which baseline freezing was recorded. Mice then received three 2-second, 0.5 mA foot shocks, each along with a 30-second auditory cue (conditioned stimulus, CS), administered at 90-second intervals. Contextual fear memory was assessed 24 hours after conditioning by returning mice to the identical chamber for 5 minutes in the absence of the CS, and percent time freezing was recorded. Cued fear memory was assessed in a novel context 24 hours after conditioning: mice were first allowed a 3-minute acclimation period in the novel chamber, followed by a 3-minute CS presentation, and percent freezing during each phase was recorded. All behavioral recordings were analyzed using Ethovision XT (Noldus).

### Immunoblotting

Tissue samples from the brain regions were lysed in cold RIPA buffer supplemented with phosphatase inhibitor and protease inhibitor cocktail tablets (Thermo Fisher Scientific, Waltham, MA, USA) using a FastPrep 24 (M.P. Biomedicals, Santa Ana, CA, USA) with bead-beating tubes (16466–042) from (VWR, Radnor, PA, USA) and zirconium ceramic oxide bulk beads from (Thermo Fisher Scientific, Waltham, MA, USA). Protein lysates were then centrifuged at 13,300 rpm for 10 min and the supernatant was collected. Protein concentration was determined by Bradford (Pierce Biotechnology, Waltham, MA, USA). 20 μg protein was separated by SDS–PAGE (sodium dodecyl sulfate–polyacrylamide gel electrophoresis) on 8%, 10%, or 16% resolving gels (ThermoFisher Scientific, Waltham, MA, USA) and transferred to PVDF membrane (EMD Millipore, Burlington, MA, USA). Autophagy markers including autophagy proteins ATG5, ATG7, and ATG16L1, as well as autophagosome formation marker light chain 3A/B (LC3A/B), and the autophagy receptor p62 (sequestosome 1, SQSTM1) were immunoblotted. Tau pathology was assessed by Western blotting with anti-tau antibody (p-Tau Thr231). Antibody vendors, catalog numbers and the dilution used are provided in **Table S1**. Imaging was performed using an Image Quant 800 (Amersham). Quantification was performed by densitometry using NIH ImageJ software.

### Histology for AD neuropathology markers

Mice were euthanized by cervical dislocation after 3 hours fast, and the right hemisphere was fixed in formalin for histology whereas the left hemisphere was snap-frozen for biochemical analysis. For amyloid plaque staining, briefly, brain sections were deparaffinized and rehydrated according to standard protocol. For epitope retrieval, mounted slides were pretreated in 70% formic acid at room temperature for 10 min. Tissue sections were subsequently blocked with normal goat serum (NGS) at room temperature for 1 hr, then incubated with monoclonal antibodies 6E10 (1:100), at 4°C overnight. Aβ immunostained profiles were visualized using diaminobenzidine chromagen. For p-Tau staining and glial activation, brains were analyzed with p-Tau Thr231, anti-GFAP (astrocytic marker), and anti-Iba1 (microglial marker) antibodies respectively. The following primary antibodies were used: phospho-Tau (Thr231) monoclonal antibody (AT180) (Thermo Fisher Scientific; # MN1040, 1:100) anti-GFAP (Thermo Fisher; # PIMA512023; 1:1,000), anti-IBA1 (Abcam; #ab178847; 1:1,000). Sections were imaged using an EVOS microscope (Thermo Fisher Scientific Inc., Waltham, MA, USA) at a magnification of 4X,10X, 20X and 40X magnification. Image-J was used for quantification by converting images into binary images via an intensity threshold and positive area was quantified.

## Statistical Analysis

All statistical analyses were conducted using Prism, version 11.0.0 (GraphPad Software Inc., San Diego, CA, USA). Comparisons between two independent groups were done using Welch’s t test and tests involving multiple factors were analyzed by either a two-way analysis of variance (ANOVA) with surgery, time or brain region as variables or by one-way ANOVA, followed by a Sidak’s post-hoc test as specified in the figure legends. Alpha was set at 5% (p < .05 considered to be significant). Data are presented as the mean ± SEM unless otherwise specified.

## Acknowledgments

We thank all members of the Harris (WiSLiM) and Lamming labs for their feedback and Drs. Galmozzi and Alexander for their insightful discussions. D.A.H. is supported by the UW Department of Surgery, School of Medicine and Public Health, Wisconsin Alumni Research Foundation, and the Office of the Vice Chancellor for Research. Additionally, D.A.H. has funding through the NIA (R03AG088813), Wisconsin Alzheimer’s Disease Research Center (P30-AG062715), and a grant from the Wisconsin Partnership Program at the UW School of Medicine and Public Health (ID 6770-2024). DAH received the Vilas Faculty Early Career Investigator Award at UW Madison. We would like to thank Medtronic (Minneapolis, MN) for a grant allowing the purchase of linear cutting staplers. RB is a UW Distinguished Research Fellow; support for this research was provided by the University of Wisconsin-Madison Office of the Vice Chancellor for Research with funding from the Wisconsin Alumni Research Foundation. The Lamming lab is supported in part by the NIA (AG056771, AG081482, AG084156, AG085898, and AG094153) and was supported by the NIDDK (DK125859), the Wisconsin Partnership Program, and startup funds from UW–Madison.

D.A.H. and D.W.L. are members of the Wisconsin Nathan Shock Center of Excellence in the Basic Biology of Aging, P30 AG092586. The Lamming lab was supported in part by the US Department of Veterans Affairs (I01-BX004031 and IS1-BX005524), and this work was supported using facilities and resources from the William S. Middleton Memorial Veterans Hospital. The content is solely the responsibility of the authors and does not necessarily represent the official views of the NIH. This work does not represent the views of the Department of Veterans Affairs or the United States Government.

## Author contributions

R.B., J.I., D.W.L., and D.A.H. conceived of and designed the experiments. R.B., J.I., M.M.S., G.Z., T.M., T.M., M.P., C.W., F.X., J.W., D.W.L., and D.A.H. analyzed the data. R.B.,and D.A.H. wrote the manuscript.

## Supplementary Figure Legends

**Supplementary Figure 1: Sleeve gastrectomy reduces adiposity and improves insulin sensitivity independent of food intake in chow-fed WT mice.** (A) Body weight of female WT mice longitudinally tracked over 12 months post-surgery. (B) Percent fat mass, (C) percent lean mass, and percent adiposity (D) assessed by body composition analysis at 6 months post-surgery. (E) Average daily food intake monitored over 12 months post-surgery. (F) Energy expenditure (EE) normalized to lean mass at 6 months post-surgery shown with ANCOVA-derived slope and intercept p-values. (G) Respiratory exchange ratio (RER) during light and dark phases at 6 months post-surgery. (H) Oral glucose tolerance test (oGTT) performed at 2 months post-surgery with corresponding area under the curve (AUC). (I) Intraperitoneal insulin tolerance test (ipITT) performed at 2 months post-surgery with corresponding AUC. (B–D, H-I) Group differences in AUC were assessed by Welch’s t-test (G) The overall effects of surgery (SG), time (T), and their interaction (T×SG) from two-way ANOVA; *p<0.05, **p<0.01 from Sidak’s multiple comparisons test. Data represented as mean ± SEM.

**Supplementary Figure 2: Energy balance in chow-fed 3xTg-AD mice 10 months post-surgery** (A) Respiratory exchange ratio (RER) and energy expenditure (EE) during light and dark phases at 10 months post-surgery in Sham and SG female 3xTg-AD mice. Statistics reflect the overall effects of surgery (SG), time (T), and their interaction (T×SG) from a two-way ANOVA. Data are represented as mean ± SEM.

**Supplementary Figure 3: Chow-fed WT mice do not exhibit cognitive deficits** (A) Latency of target in Barnes Maze acquisition period over the five days of training and in short term memory (STM) and long-term memory (LTM) tests at 7 months in WT mice. (B) The number of error hole visits during Barnes maze training phase in STM and LTM tests at 7 months in WT mice. (C) Percent inactivity recorded during the cued test phase of the fear conditioning paradigm at 7 months in WT mice. (D) Percent inactivity recorded during the acclimation, conditioning, context, and cued phases of the fear conditioning paradigm at 7 months in WT mice (E) The preference for a novel object over a familiar object was assayed at 7 months via STM and LTM tests. The dashed lines at +0.2 and -0.2 indicates the threshold for discrimination index (DI) values showing the preference for novel or familiar objects. (A-E) Statistics for the overall effects of surgery, time and the interaction represent the p value from a 2-way ANOVA. * p<0.05 **p<0.01, ****p<0.0001 from Sidak’s multiple comparison test.

**Supplementary Figure 4. Glucose homeostasis is comparable between SG and Sham mice at 9 months post-surgery in WD-fed 3xTg-AD mice.** (A) Oral glucose tolerance test (oGTT) and corresponding area under the curve (AUC) at 9 months post-surgery in Sham-WD and SG-WD female 3xTg-AD mice. (B) Intraperitoneal insulin tolerance test (IpITT) and corresponding AUC at 9 months post-surgery. ns = not significant. Data are represented as mean ± SEM.

## Supplementary Table Legends

**Supplementary Table 1:** Antibodies used for both western blotting and immunohistochemistry

