## Supplementary figures and images for "Sleeve gastrectomy improves metabolic health, cognition, and Alzheimer’s Disease pathology in 3xTG mice"

### Supplemental Figure 1

#### Supplementary Figure 1

WT

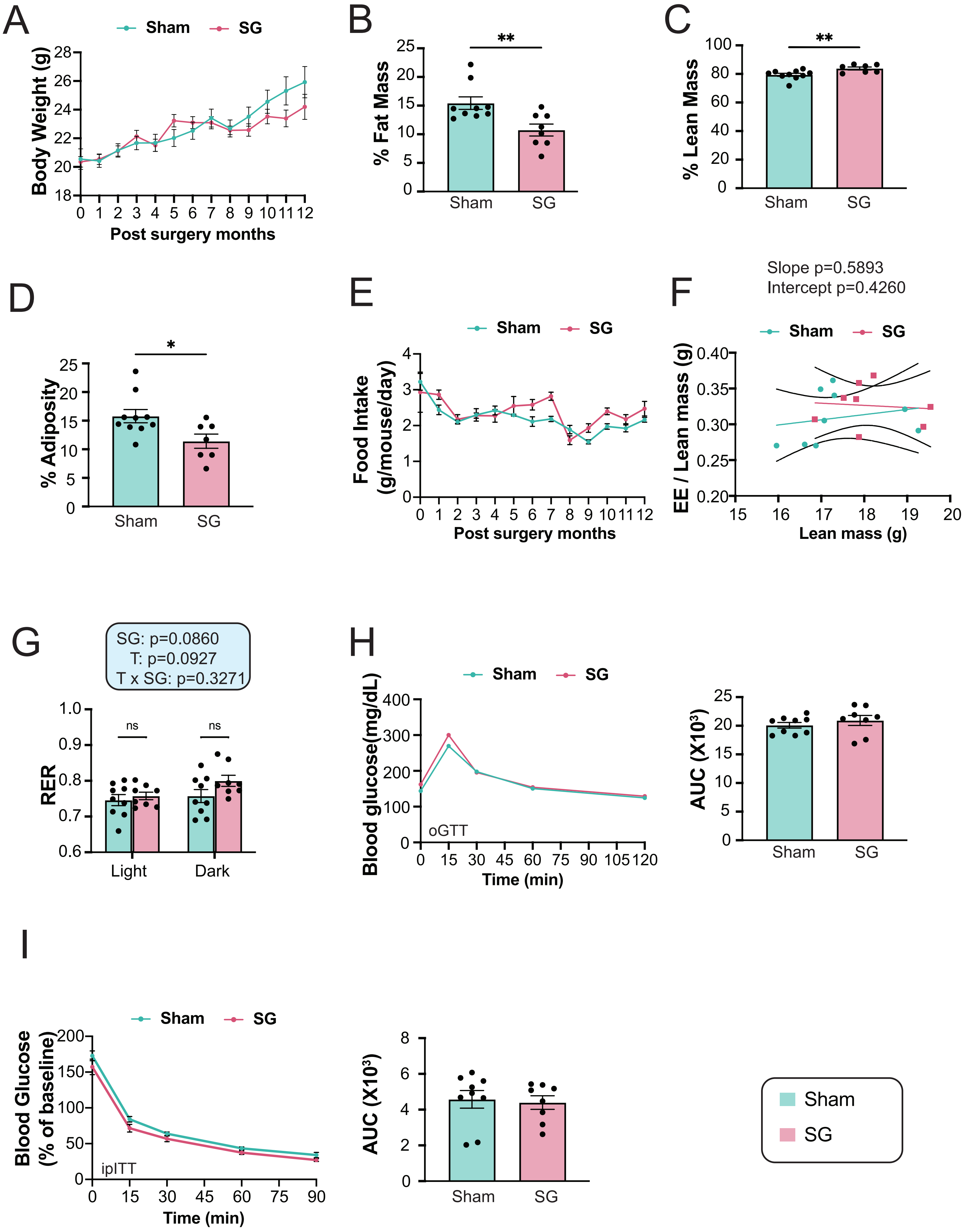

### Supplemental Figure 2

#### Supplementary Figure 2

3xTg

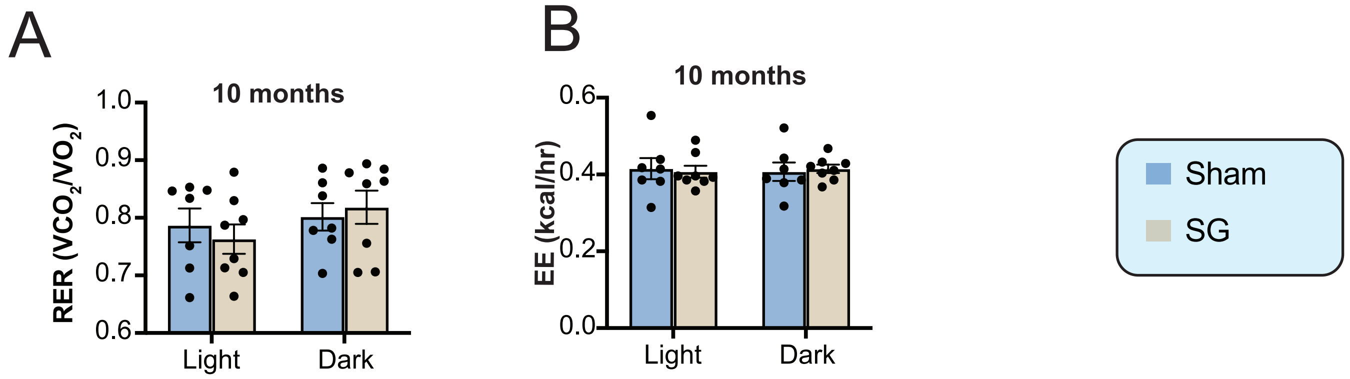

### Supplemental Figure 3

#### Supplementary Figure 3

WT

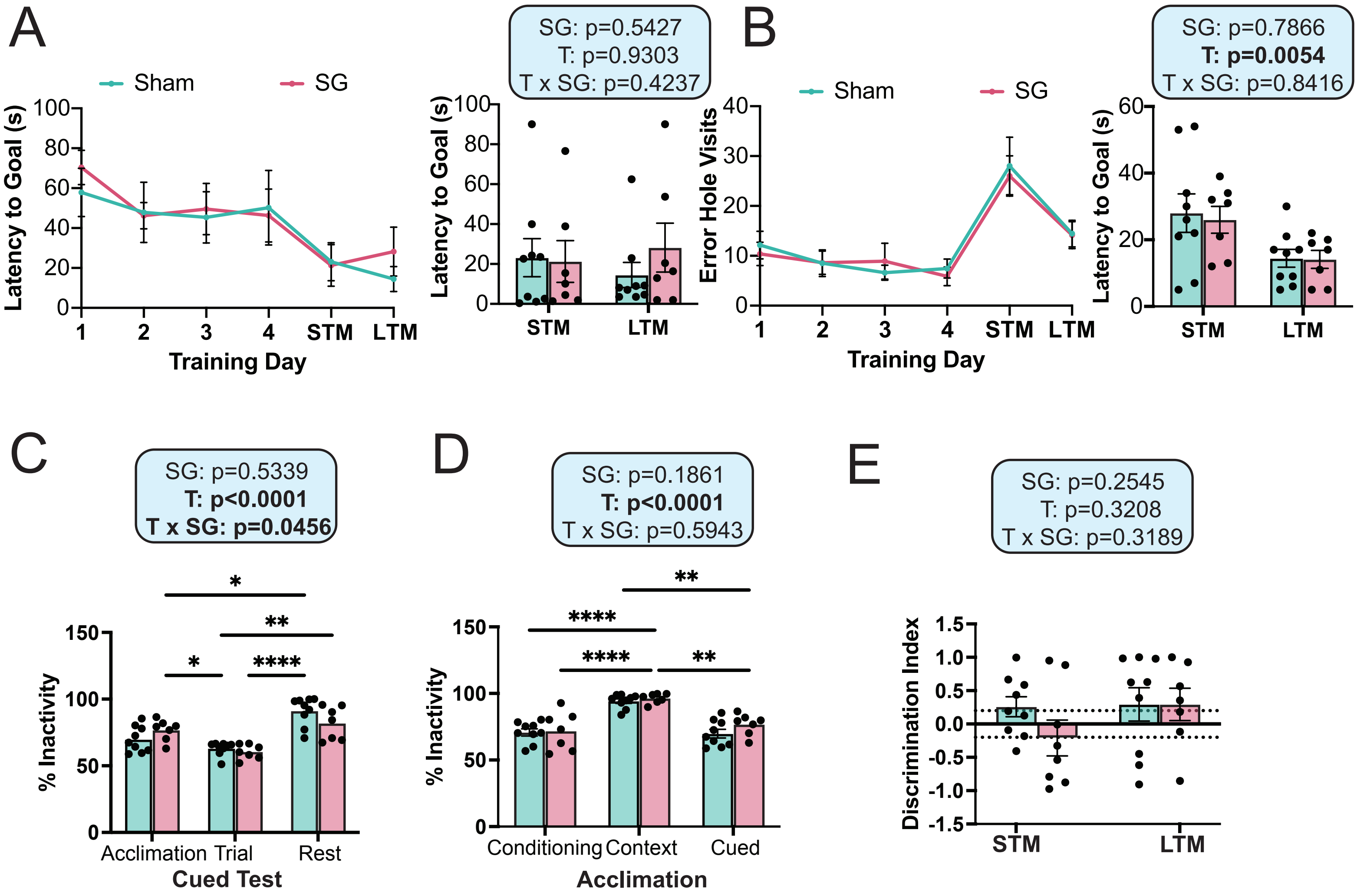

### Supplemental Figure 4

#### Supplementary Figure 4

3xTg-WD

A

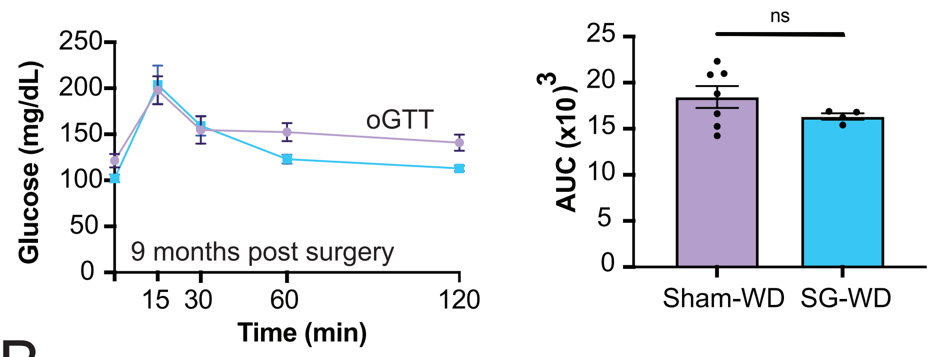


B

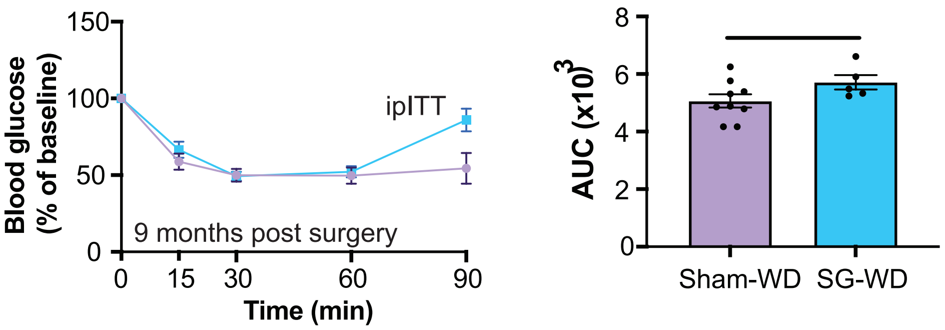
